# Maize inbred line B96 is the source of large-effect loci for resistance to generalist but not specialist spider mites

**DOI:** 10.1101/2021.02.04.429847

**Authors:** Huyen Bui, Robert Greenhalgh, Gunbharpur S. Gill, Meiyuan Ji, Andre H. Kurlovs, Christian Ronnow, Sarah Lee, Ricardo A. Ramirez, Richard M. Clark

**Affiliations:** School of Biological Sciences, University of Utah, 257 South 1400 East, Salt Lake City, Utah 84112, USA; Department of Biology, Utah State University, 5305 Old Main Hill, Logan, Utah 84322, USA; Henry Eyring Center for Cell and Genome Science, University of Utah, 257 South 1400 East, Salt Lake City, Utah 84112, USA

**Keywords:** Two-spotted spider mite, *Tetranychus cinnabarinus*, Banks grass mite, bulked segregant analysis, antibiosis

## Abstract

Maize (*Zea mays* subsp. *mays*) yield loss from arthropod herbivory is substantial. While the basis of resistance to major insect herbivores has been comparatively well-studied in maize, less is known about resistance to spider mite herbivores, which are distantly related to insects and feed by a different mechanism. Two spider mites, the generalist *Tetranychus urticae*, and the grass-specialist *Oligonychus pratensis*, are notable pests of maize, especially during drought conditions. We assessed the resistance to both mite species of 38 highly diverse maize lines, including several previously reported to be resistant to one or the other mite species. We found that line B96, as well as its derivatives B49 and B75, were highly resistant to *T. urticae*. In contrast, neither these three lines, nor any others included in our study, were notably resistant to *O. pratensis*. Quantitative trait locus (QTL) mapping with F2 populations from crosses of B49, B75, and B96 to susceptible B73 identified a large-effect QTL on chromosome 6 as underlying *T. urticae* resistance in each line, with an additional QTL on chromosome 1 in B96. Genome sequencing and haplotype analyses identified B96 as the apparent sole source of resistance haplotypes. Our study identifies loci for use in maize breeding programs for *T. urticae* resistance, as well as to assess if the molecular-genetic basis of spider mite resistance is shared with insect pests of maize, as B96 is also among the most resistant known maize lines to several insects, including the European corn borer, *Ostrinia nubilalis*.

**Key message Maize:** (*Zea mays* subsp. *mays*) inbred lines B49, B75, and B96 harbor large-effect loci for resistance to the generalist spider mite *Tetranychus urticae*, but not the specialist *Oligonychus pratensis*.

## Introduction

While cereal crops including maize (*Zea mays* subsp. *mays*) provide more than half of human calories, yield losses to both abiotic and biotic factors are substantial, and can be synergistic (Deutsch et al. 2018). Among abiotic factors, drought stress, which is often associated with elevated temperatures, is especially important, and is projected to become more so in many regions under current climate change projections (Snowdon et al. 2020). Further, in spite of extensive pesticide use, as much as ∼20% of global maize production is lost due to herbivory by arthropod pests (Oerke 2006). Taking into account anticipated climate change impacts on herbivores, which include increased metabolic rates and winter survival, yield losses from herbivory by insects for maize may increase as much as ∼30% given an average global surface temperature increase of 2°C (Deutsch et al. 2018).

Grasses are attacked by insect herbivores of diverse feeding guilds, such as leaf chewing (e.g., caterpillars and grasshoppers), stem mining (e.g., corn borers), and phloem-feeding (e.g., aphids) (Meihls et al. 2012). Additionally, spider mites (Arthropoda: Chelicerata: Arachnida: Acari), which belong to a sister taxon to insects, have long been recognized as field pests of maize and related cereals (Owens et al. 1976). As opposed to the major insect herbivores of maize, spider mites feed on individual leaf mesophyll cells with specialize mouth parts (needle-like stylets) (Bensoussan et al. 2016; Bui et al. 2018). Although only ∼0.6 mm in length, a single spider mite female can lay dozens of eggs, and the generation time can be as little as seven days at high temperatures (Owens et al. 1976; Grbic et al. 2007). Population sizes on single plants may therefore reach tens of thousands during a growing season, and can result in nearly complete yield loss (Owens et al. 1976).

Two spider mite species, the generalist two-spotted spider mite (*Tetranychus urticae*), and the grass-specialist Banks grass mite (*Oligonychus pratensis*), are significant pests on maize (Owens et al. 1976; Archer and Bynum 1993; Peairs 2010). While the impact of these herbivores on maize has been most studied in the USA, *T. urticae* is globally distributed, and *O. pratensis* has been reported on multiple continents (Migeon et al. 2010). In arid regions of the Western USA, both species can be economically damaging field pests of maize. In particular, yield losses from spider mite herbivory typically occur during hot and dry conditions as a result of rapid mite generation times at high temperatures, altered relationships with natural pathogens or predators, or potentially changes in the physiology of drought-stressed plants that favor the herbivore (Peairs 2010; Gill et al. 2020). In less arid regions of Eastern North America, *T. urticae* is the principal spider mite pest of maize, and causes little damage in most years. However, during periods of abnormally high heat and low precipitation, populations of *T. urticae* can increase rapidly, and this species can become a dominant arthropod pest of maize, as occurred during the severe drought of 2012 in states like Iowa, USA (Al-Kaisi et al. 2013). Such outbreaks are very difficult to control, as spider mites, especially the generalist *T. urticae*, have among the highest known rates of pesticide resistance evolution (Van Leeuwen et al. 2010).

The inability to effectively control spider mites with pesticides, coupled with their importance as pests during hot and dry conditions, has generated interest in understanding the extent and magnitude of spider mite resistance in maize germplasm, and variation in resistance has been observed for both *T. urticae* and *O. pratensis* in multiple studies (Owens et al. 1976; Archer 1987; Kamali et al. 1989; Mansour, F. and Bar-Zur, A. 1992; Mansour et al. 1993; Tadmor et al. 1999; Bynum et al. 2004a, b). It should be noted that some of these older studies reference *T. cinnabarinus*, which is now thought to be the red color form of *T. urticae* (Auger et al. 2013). Previous efforts to assess maize resistance have varied in several ways, such as in the stage of maize plants used for screening, in whether whole plants or detached leaves were employed, in the genetic composition of both the maize lines and mite strains, and in how resistance was measured. With respect to the latter, resistance arising from antibiosis (impaired herbivore growth or reproduction), or from tolerance (the ability of a plant to support high herbivore populations without detrimental effects), has been assessed. For *T. urticae*, several maize lines, including Oh43 and B96 (formerly called 41.2504B), have been reported to exhibit strong antibiosis (Kamali et al. 1989; Mansour, F. and Bar-Zur, A. 1992; Tadmor et al. 1999). Resistance arising from antibiosis, but also tolerance, has been reported for *O. pratensis* (Bynum et al. 2004b), or alternatively the mechanism of resistance was not determined (Owens et al. 1976).

Despite variation in maize resistance to both mite species, the underlying molecular-genetic basis is largely unknown, as is whether resistance mechanisms are shared with insect herbivores. To begin to address these questions, Bui et al. (2018) examined transcriptional responses to spider mites using B73, a comparatively spider mite susceptible maize line that is the source of the reference maize genome. B73 responses to both *T. urticae* and *O. pratensis* resembled that of mechanical wounding (tissue disruption is a component of feeding by all herbivores), and were similar to responses reported for herbivory by *Spodoptera exigua*, a lepidopteran caterpillar (Tzin et al. 2017; Bui et al. 2018). In particular, genes associated with jasmonic acid, a phytohormone known to mediate plant responses to many herbivores (Howe and Jander 2008), were upregulated. Moreover, genes for the synthesis of benzoxazinoids, like 2,4-dihydroxy-7-methoxy-1,4-benzoxazin-3-one (DIMBOA), and its derivative compounds, were upregulated by both mite species. Benzoxazinoids are defensive compounds that are widespread in grasses (family Poaceae) and are stored as glucoside conjugates; upon tissue disruption during feeding, the aglucones are generated, and have detrimental effects on susceptible herbivores, including many lepidopteran larvae (Wouters et al. 2016). In maize, benzoxazinoids are synthesized by 14 enzymes encoded by the *BX1*-*BX14* genes. Using maize *bx1* and *bx2* mutants that lack these compounds, Bui et al. (2018) found that on maize seedlings reproduction of *T. urticae*, but not *O. pratensis*, increased in the absence of benzoxazinoids. Although this finding suggests that some aspects of maize resistance to insects are shared with spider mites, whether variation in benzoxazinoid content among maize lines accounts for natural variation in resistance to *T. urticae* is unknown (Bui et al. 2018), as is the basis of resistance to *O. pratensis*.

To address these outstanding questions, we surveyed 38 inbred lines for antibiosis to both *T. urticae* and *O. pratensis*. We included lines previously reported to be resistant to spider mites, as well as lines reported to be resistant to lepidopteran herbivores, or to have high DIMBOA content. Further, to facilitate follow up genetic studies, we included the diverse founders of the maize Nested Association Mapping (NAM) population, a high-resolution genetic mapping resource (McMullen et al. 2009). We identified only a small number of maize lines as highly resistant to *T. urticae*, and no lines in our study exhibited strong antibiosis to *O. pratensis*. QTL mapping using three highly *T. urticae* resistant lines revealed two prominent loci as underpinning antibiosis to *T. urticae*. Our findings inform studies of plant resistance to spider mite herbivores of varying host breadth, and identify loci for use in breeding programs – as well as candidate genes – for resistance to *T. urticae* in maize.

## Materials and methods

### Selection of maize lines and crosses

Seeds for maize inbred B73 were kindly provided by G. Drews (University of Utah, Salt Lake City, UT, USA), while those for 37 other maize inbred lines were obtained from the North Central Regional Plant Introduction Station (Ames, IA, USA). Apart from B73, these included 24 additional parents of the NAM recombinant inbred line population (McMullen et al. 2009). We failed to propagate one NAM line founder, Ki3, which was therefore excluded from our study. The remaining 13 maize lines were selected because they met one or more of the following criteria: (1) were previously reported to have moderate to high resistance to either *T. urticae* or *O. pratensis* (B49, B64, B96, Oh43, Tx202, and TAM-MITE1, with TAM-MITE1 reported to exhibit antibiosis to both mite species; Oh43 is also a NAM line founder) (Kamali et al. 1989; Mansour, F. and Bar-Zur, A. 1992; Tadmor et al. 1999; Bynum et al. 2004b; and phenotypic data as reported in the U.S. National Plant Germplasm System, GRIN); (2) were parents (B14 and B52) or resultant lines (B49, B64, B68, and B86) in breeding programs involving crosses with B96 or Oh43, which have been reported to be highly *T. urticae* resistant (pedigree data is available at the Maize Genetics and Genomics Database, or MaizeGDB) (Portwood et al. 2019); (3) are widely used maize inbred lines with extensive genetic and genomic resources (Mo17 and W22) (Portwood et al. 2019); (4) have been reported to be resistant to *Ostrinia nubilalis*, the European corn borer, in previous studies or as referenced in MaizeGDB (B49, B64, B52, B75, B86, B96, and CI31A) (Kamali et al. 1989; Barry et al. 1994; Portwood et al. 2019); or (5) have been reported to have high levels of DIMBOA in previous studies, or are labelled with the phenotype descriptor “DIMBOA content high” in MaizeGDB (B75, B96, and CI31A) (Bing et al. 1990b; Barry et al. 1994; Portwood et al. 2019). Crosses of B73 to B49, B75, and B96 were performed in greenhouses at the University of Utah, and F2 seeds for mapping studies were subsequently produced by selfing F1 progeny.

### Spider mite stocks and propagation

A *T. urticae* strain, W-GR, and an *O. pratensis* strain, were maintained at high population sizes (several thousand mites) on B73 maize plants in isolated laboratory rooms as previously described at the University of Utah and Utah State University (Logan, UT, USA) (Bui et al. 2018). The origin of these strains, neither of which is inbred, was described by Bui et al. (2018). Briefly, WG-R was derived from *T. urticae* mites collected at several sites, including from maize plants in a greenhouse, and the *O. pratensis* strain was isolated from field-grown maize. Both stocks originated from Utah, USA. We also used five additional strains of *T. urticae* collected in Utah, USA: Catnip (collected from a catnip plant, *Nepeta cataria*), NightS (collected from a bittersweet nightshade plant, *Solanum dulcamara*), ShCo (collected from a morning glory plant, *Ipomoea purpurea*) and strains KH and WGDel (collected from populations on multiple adjacent plant species, not including maize; KH was collected from house plants and WGDel was collected from a greenhouse). These five strains were inbred by mother-son crosses for five generations (Catnip, KH and ShCo) or eight generations (NightS and WGDel) on detached common bean (*Phaseolus vulgaris*) leaves maintained on wet cotton as previously described (Bryon et al. 2017). While two of these inbred strains – ShCo and WGDel – were reported previously (Wybouw et al. 2019), the inbred Catnip, KH, and NightS strains were produced as part of the current work. Unless otherwise noted, the five inbred *T. urticae* strains were maintained by serial passage on detached bean leaves at the University of Utah as described previously (Bryon et al. 2017).

### Screen of 38 maize lines for spider mite resistance

Seeds for the 38 maize inbred lines were sown 1-2 cm deep in 5 × 5 cm plastic pots, and resulting seedlings were transplanted to 20 cm diameter pots 10 days after sowing. Germination and subsequent propagation at University of Utah greenhouses used a 16h-light/8h-dark photoperiod, approximate temperature of 25°C, and Metro-Mix® 900 growing mix soil (Sun Gro® Horticulture, Fillmore, UT, USA). After transplanting, plants were kept in trays and watered from below; the plants were fertilized weekly with 200 ppm NutriCulture Cal-Mag Special 16N-3P-16K (Plant Marvel Laboratories, Chicago Heights, IL, USA). At eight weeks, barriers were applied on the 8^th^ leaf using Tanglefoot, a non-phytotoxic wax (The Scotts Miracle-Gro Company, Marysville, OH, USA) as previously described (Bui et al. 2018; Gill et al. 2020). For each plant, three barriers were established perpendicular to the length of the leaf blades to define two adjacent enclosures of 6.5 cm in length. For each maize inbred line and each spider mite species, three plants were used, resulting in a total of six enclosures.

Twenty-four hours after enclosure establishment, 1-to 2-day-old adult females of *T. urticae* (strain W-GR) or *O. pratensis* were introduced into each enclosure. Briefly, the females were collected from mite populations synchronized on detached B73 maize leaves as described by Bui et al. (2018). Ten *O. pratensis* or eight *T. urticae* W-GR females were sucked into barrier pipet tips by vacuum (given the small size of mites, and their density on leaves, occasionally several additional mites were captured in tips, but see “effective female” correction below). The mites were then tapped to the bottom of the tips against the barriers, the tops of the tips were cut off to allow mites to escape, and tips were taped to the underside of the leaves between two Tanglefoot barriers (one tip per enclosure) as previously described (Bui et al. 2018; Gill et al. 2020). It takes a few hours for mites to exit tips and start feeding, and some may fail to exit tips, or fall off while exiting. Therefore, the number of females in each enclosure was assessed by visual inspection one day after release, as well as at the end of the experiment (at six days). The number was assessed twice for reproducibility, and to ensure that a mite was not overlooked at day one by visual inspection, which is a challenge given spider mites’ small size. The greater of these counts was considered to be the number of females that had successfully entered a given enclosure (hereafter effective females). Six days after adding tips with mites to enclosures, individual enclosures were collected by cutting immediately adjacent to Tanglefoot barriers, and the resulting leaf sections were transferred to 4°C (which arrests mite reproduction and development). The total number of progeny (eggs and viable mites, all stages) in each enclosure was then determined under a dissecting microscope and normalized on a per enclosure basis by the count of effective females.

The scope of screening 38 maize lines for resistance against two spider mite species imposed several constrains on the experimental design. Screening was performed in two batches (batches 1 and 2, see Fig. 1); moreover, for logistical reasons we did not use a randomized design for the survey of the 38 lines (although within each batch, resistance to both mite species was assessed simultaneously for each maize line). To allow inter-batch reproducibility to be assessed, however, four maize lines were included in both batches (Fig. 1 and Results).

**Fig. 1.**
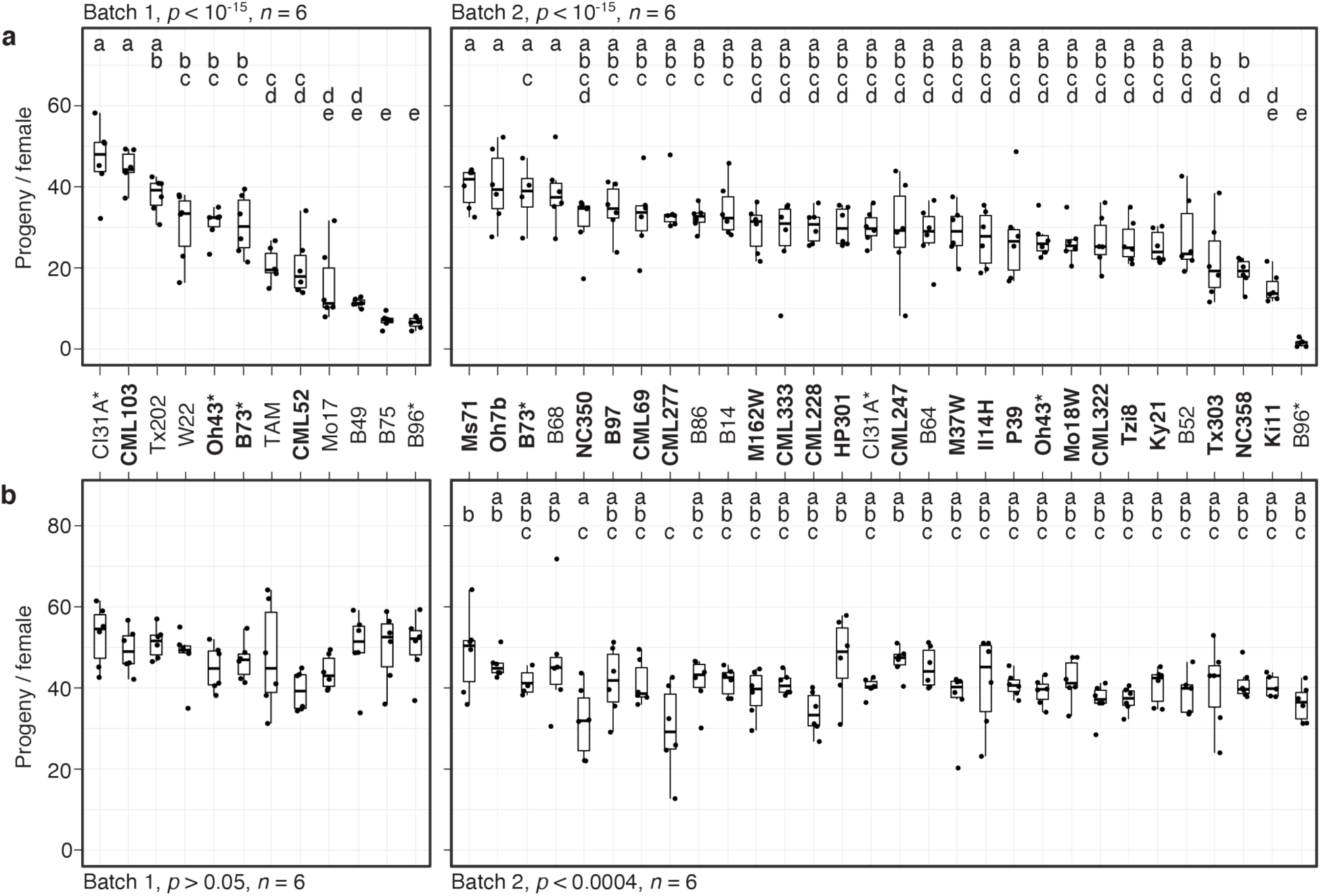
Variation in resistance among 38 maize inbred lines to *T. urticae* and *O. pratensis*. Indicated is the number of progeny per adult *T. urticae* female (**a**) and *O. pratensis* female (**b**) by maize line after six days of feeding in enclosures on the 8^th^ leaves (boxplots with overlay of data points are shown). The screen was performed in two batches, 1 and 2 (left and right in each panel, respectively). In panel **a**, maize lines are ranked by median resistance – least to most, left to right –within each batch; to facilitate comparisons between mite species, the same ranking is used in panel **b**. Where significant ANOVA results (*p* < 0.05) were obtained by batch, pairwise significance was assessed with correction for multiple testing using the Hochberg method (different letters denote significant differences at *p* < 0.05). Maize line founders of the NAM population are in bold, and four lines included in both batches are indicated by asterisks. Sample size, *n* (enclosures per maize line, see Materials and Methods), are as indicated except for maize lines B73 (**a** and **b**), B52 (**b**), or Ki11 (**b**) in batch 2, where either four or five enclosures were successfully established. TAM: TAM-MITE1.

### Validation of spider mite resistance

Spider mite resistance for lines B49, B73, B75, and B96 was subsequently assessed at a second site (a Utah State University greenhouse) using four replicates per maize line per mite species in a completely randomized design. Seeds were sown in 22.5 cm pots using Sunshine 3 soil (Sun Gro® Horticulture, Fillmore, UT, USA). Each pot was automatically fertigated (irrigation + fertilizer) at the rate of 0.3 L/day using 21-5-20 water soluble fertilizer mixture (4.8 kg/100L of water, Peters Excel supplied by ICL Specialty Fertilizers, Summerville, SC, USA). Plants were kept at approximately 25°C with a 16h-light/8h-dark photoperiod cycle. When plants were 8-weeks old, Tanglefoot enclosures of length 15 cm were established on the 8^th^ and 9^th^ leaves (the enclosures were centered on leaf midpoints). Twenty adult female mites from either the *T. urticae* W-GR strain or the *O. pratensis* strain collected from whole B73 plants (see Material and Methods section “Spider mite stocks and propagation”) were then introduced into the enclosures. Mite collection into barrier tips, and placement on leaves, was as for the initial survey of the 38 lines. After six days of mite infestation, the leaf samples inside each Tanglefoot enclosure were collected and frozen (to stop mite development and reproduction). The number of progeny (eggs and mites of any stage) in each enclosure were subsequently counted after leaf samples were thawed; progeny per female was then calculated on a per plant basis (the sum of the progeny in enclosures on the 8^th^ and 9^th^ leaves were divided by 40).

### Performance of five inbred *T. urticae* strains on maize inbreds

The productivity of five additional *T. urticae* strains (inbred strains Catnip, KH, NightS, ShCo, and WGDel) on maize lines B49, B73, B75, and B96 was assessed in greenhouse bays at the University of Utah. To allow a comparison to earlier findings, the *T. urticae* W-GR strain was also included. Prior to collecting females for use in resistance screening, mites from the five inbred strains, for which stocks were maintained on detached bean leaves, were passaged for ∼2 generations on detached B73 maize leaves for physiological adaptation. Subsequently, the experimental design was identical to that used for the initial survey of the 38 maize lines, except that single enclosures per plant were established on the 8^th^ leaves, six replicates (single plants) were used per *T. urticae* strain and maize inbred line, and a completely randomized design was employed.

### Design for resistance mapping, phenotyping, and sample preparation

To localize maize resistance loci to *T. urticae* in F2 populations derived from B49 × B73, B75 × B73, and B96 × B73 crosses, we used bulked segregant analysis (BSA) genetic mapping (Zou et al. 2016; Kurlovs et al. 2019). For phenotyping, 400 plants from each F2 population were sown to evaluate *T. urticae* resistance; for logistical reasons, experiments for each cross were performed with two subsets of 200 plants each (hereafter termed replicates) grown in greenhouse bays at the University of Utah. For each replicate of each cross, seeds were sown in 5 × 5 cm plastic pots with Metro-Mix 900 growing mix soil. Ten days after sowing, the seedlings were transplanted into 20 cm diameter pots and watered and fertilized as described previously. Tanglefoot enclosures of 6.25 cm in length were established on the 4^th^ leaves of 3-week-old plants. Forty adult females of the *T. urticae* W-GR strain, collected from B73 plants, were released into each enclosure using barrier pipet tips as already described. Six days after infestation, the extent of mite feeding damage in each enclosure was scored using a visual scale (1 for the least damage, 7 for the heaviest damage). Each plant was scored by at least two project participants to produce an average score.

If, in a given enclosure, more than 10 mites (25% of the released mites) were observed to have been trapped in a pipet tip, the sample was excluded from further analyses. Subsequently, damage scores were sorted to identify the 50 most sensitive plants (heavier damage, higher score) and 50 most resistant plants (lesser damage, lower score). For DNA preparation, single 4 mm leaf punches from all plants in a given pool were collected, combined, and DNA was extracted using the DNeasy PowerPlant Pro kit (Qiagen, Germantown, MD, USA) following the manufacturer’s recommendations. Illumina DNA-seq libraries of 550 bp average insert sizes were then constructed at the High Throughput Genomics Core Facility at the University of Utah and 125-bp paired-end DNA reads were generated on an Illumina HiSeq 2500 sequencer to generate between 26.8 and 59.3 million read pairs per sample (∼2.8 to 6.2-fold coverage depth for the maize genome).

### Read alignment, variant calling and BSA scans

We used QTLseqr (version 0.7.2) (Mansfeld and Grumet 2018), which implements the G’ method (Magwene et al. 2011), to detect differences in allele frequencies between pools that, as assessed with a false discovery rate (FDR) estimation, identify QTL regions. The input for the G’ method is genotypic information at single nucleotide polymorphism (SNP) sites as inferred from respective allele depths of aligned sequence reads. Reads from pooled samples were aligned to the B73 maize reference genome version AGPv4 (Jiao et al. 2017) using the default settings of BWA (version 0.7.15-r1140) (Li 2013), and sorted by position using SAMtools 1.3.1 (Li et al. 2009). SNP detection was performed with the Genome Analysis Toolkit (GATK) version 3.6-0-g89b7209 (McKenna et al. 2010); in accordance with GATK best practices recommendations (Van der Auwera et al. 2013), duplicate reads were marked using Picard Tools 2.6.0-SNAPSHOT (http://broadinstitute.github.io/picard) prior to indel realignment with GATK. As QTLseqr only allows the specification of two samples for a given analysis (“HighBulk” and “LowBulk”, respectively), SAMtools was used to merge the resistant and sensitive replicates for analyses using both replicates by maize line. SNPs were called using the GATK UnifiedGenotyper tool for each bulked-segregant experiment, with SNPs for the individual and combined replicates called separately for each cross. The GATK VariantsToTable tool was used to generate tab-delimited table files (with the following fields: CHROM, POS, REF, ALT, AD, DP, GQ, and PL) from each of the respective UnifiedGenotyper variant call format (VCF) files to produce the required input for QTLseqr.

For each of the six resultant input table files by replicate (i.e., two replicates for each of the three crosses), reads were filtered by adapting QTLseqr recommendations (Mansfeld and Grumet 2018). Briefly, depending on sequencing depth by sample, the following SNP read coverage requirements were used: a minimum depth across both pools in a replicate of 6-8, a maximum depth of 24-36, a maximum depth difference between the resistant and sensitive pools of 6-8, and minimum sample depths of 3-4 (these parameters resulted in filtered SNP counts of between 1.90 to 4.78 million among replicates across the three crosses). In combining replicates by cross, the respective parameters were doubled, resulting in filtered SNP counts by cross of between 3.15 to 6.58 million. Further, in all cases, a reference allele frequency cutoff of 0.2 and a minimum genotype quality of 20 were used. Following SNP filtering, G’ analyses with window sizes of 10 million bp were used, with outliers filtered using Hampel’s method, to identify QTL regions (Mansfeld and Grumet 2018). In all cases, a false discovery rate (FDR) of 0.01 was applied to identify QTL.

### Haplotype analyses

To assess haplotype similarities between B49, B75, and B96, we generated between 255.2 to 319.2 million Illumina 125-bp paired-end reads for these three maize inbred lines (∼26.6 to 33.2-fold coverage depth), aligned the resulting reads to the B73 genome, and performed variant detection (the same methods used for the BSA analyses were employed). Regions of haplotype similarity between B49, B75, and B96 were then detected in pairwise comparisons of SNPs using 5 Mb sliding windows with 500 kb offsets. For this analysis, 5.69 million high-confidence SNPs identified across all three lines were selected with a filtering scheme modified from GATK documentation specifying guidelines for hard-filtering. SNPs included in the analysis had to have: (1) read coverage depth (as described by the AD field in the variant call format v. 4.2) between 0.25-to 1.5-fold of the genome wide average (to select against copy variable regions), (2) quality score normalized by allele coverage depth (QD field) of at least two, (3) maximum strand odds ratio (SOR) of three, (4) minimum mean mapping quality score (MQ) of at least 50, (5) minimum mapping quality rank sum score (MQRankSum) of -8, and (6) rank sum test for relative positioning of reference versus alternative alleles within reads (ReadPosRankSumTest) of at least -8. For each sliding window haplotype comparison between pairs of lines, 20% of SNPs had to pass the above-listed quality control criteria in the genomic window in a given pair of lines, and at least 21 SNP positions had to present.

### QTL confirmation by single locus genotyping

A QTL interval for *T. urticae* resistance was characterized by genotyping F2 individuals for which phenotypic data was available (BSA replicate two for cross B49 × B73, and replicate one for cross B75 × B73). For genotyping, a co-dominate PCR marker at 73.4 Mb on chromosome 6 that distinguished a small indel between B73 as compared to both B49 and B75 was selected by inspection of Illumina read data. DNA was isolated from 2 mm leaf punches from frozen leaf tissue collected from individual plants using the Extracta DNA Prep kit and protocol (Quantabio, Beverly, MA, USA). PCR was performed with forward (5’-GCAGCCAGCAAGAAGAAGTCC-3’) and reverse (5’-CACAGGTCGTAGTTAGTATTCC-3’) primers using Taq polymerase, dNTPs and standard buffer (New England BioLabs, Ipswich, MA, USA) with 35 cycles of 95°C for 30 seconds, 52°C for 30 seconds, and 68°C for 60 seconds. Genotypes were assessed following resolution of PCR products on 4% agarose gels stained with ethidium bromide. The use of frozen tissue with the Quantabio DNA isolation method did not lead to successful amplification from all samples, and where amplified bands were faint (especially for heterozygotes), genotype call were not attempted (genotype calls were performed blindly to knowledge of the plant phenotype, however).

### Statistical analyses

Statistical analyses of phenotypic data were performed by analysis of variance (ANOVA) or regression (see Results) using the R language (R Core Team 2016). Where ANOVA analyses were significant, pair-wise t-tests were performed with correction for multiple testing using the Hochberg method. Where boxplots were used for data visualization, the jitter option in the R package ggplot2 (Wickham 2016) was employed (a small unit of random variation was added per data point to avoid the plotting of overlapping points). Display items produced with R, or those output from the QTLseqr program, were adjusted using Adobe Illustrator (Adobe, San Jose, CA, USA).

## Results

### Variation in mite resistance in maize

We evaluated mite resistance by surveying the number of progeny per female produced on enclosures on the 8^th^ leaves of 38 inbred maize lines that included founders of the maize NAM population (McMullen et al. 2009), as well as lines previously reported to be resistant to *T. urticae, O. pratensis*, or both (NAM founders are indicated in bold in Fig. 1; see Materials and Methods). Given the scope of the screen, it was performed in two batches (batches 1 and 2), and four lines were included in both batches to assess reproducibility (B73, B96, CI31A, and Oh43). As assessed with *T. urticae* strain W-GR, significant variation in resistance was observed in both batches (ANOVA, *p* < 10^−15^ for each batch, Fig. 1a; for display by batch, lines are ordered from least to most resistant). In each batch, B96 was the most resistant maize line as assessed by median progeny per female, and in general progeny per female for lines replicated in both batches were similar. In batch 1, three other lines, B75, B49, and Mo17, were not significantly different from B96 (t-tests, *p* < 0.05, Hochberg correction for multiple testing). Apart from B96 and B49, resistance levels to W-GR for other maize lines previously reported to exhibit antibiosis to *T. urticae*, including Oh43 and TAM-MITE1 (see Materials and Methods), were generally representative of the majority of the 38 lines tested. In contrast to the findings for *T. urticae*, we did not observe marked variation in resistance to *O. pratensis* (an ANOVA was only significant for batch 2, *p* < 0.0004), and median values for *O. pratensis* progeny per female only ranged from ∼30 to 50, including for the inbred lines that exhibited the highest levels of resistance to *T. urticae* (i.e., B49, B75, and B96, compare Fig. 1a to Fig. 1b).

For follow-up studies, we focused on the three most *T. urticae* resistant lines, B49, B75, and B96, as defined by lowest median values of progeny per female (Fig. 1a). To validate the resistance profiles for these lines, along with susceptible B73 (Fig. 1a, batches 1 and 2), we assessed resistance to both *T. urticae* and *O. pratensis* at a second greenhouse location. As compared to the initial screen of 38 lines, we used larger leaf enclosures with more mites, which were added to both the 8^th^ and 9^th^ leaves of individual plants in a completely randomized design. Overall, progeny per female for each line was lower than observed in the initial screen of the 38 lines, potentially reflecting environmental variation between locations, as well as differences in methods (compare Fig. 2 to Fig. 1, and see Materials and Methods). Nevertheless, relative resistance among the four maize lines was identical to that observed in the initial survey. Briefly, for *T. urticae*, significant variation in resistance was observed (ANOVA, *p* < 10^−4^); while B49, B75, and B96 were not significantly different from each other, all were significantly more resistant than B73 (t-tests, *p* < 0.05, Hochberg correction for multiple testing; Fig. 2a). In contrast, no significant variation in resistance to *O. pratensis* was detected (ANOVA, *p* > 0.05; Fig. 2b).

**Fig. 2.**
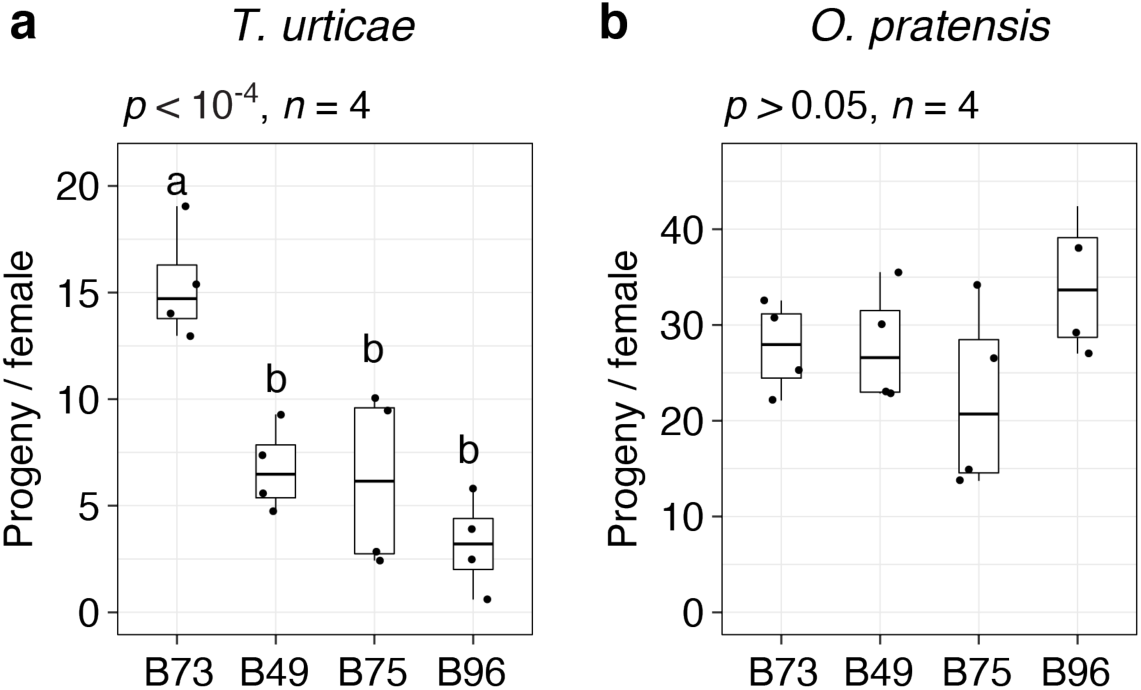
Spider mite resistance profiles for B73, B49, B75, and B96. Indicated is the number of progeny per adult *T. urticae* female (**a**) and *O. pratensis* female (**b**) for each of the four maize lines after six days of feeding in enclosures on the 8^th^ and 9^th^ leaves (boxplots with overlay of data points are shown; sample size, *n*, refers to the number of plants used). Where significant variation was observed by ANOVA (*p* < 0.05), pairwise significance was assessed with correction for multiple testing using the Hochberg method (different letters denote significant differences at *p* < 0.05).

### B49, B75, and B96 are resistant to multiple *T. urticae* strains

Prior to the current study, the outbred W-GR strain of *T. urticae* had been maintained on B73 maize plants for at least 15 generations. Host adaptation over a modest number of generations has been documented for *T. urticae* populations (Sousa et al. 2019), and it is possible that adaptation of W-GR to B73 might have impacted its relative performance on other maize lines, including B49, B75, and B96, as compared to non-adapted *T. urticae* strains. To assess the generality of our findings with the *T. urticae* W-GR strain, we therefore determined resistance for B49, B75, and B96, along with susceptible B73, to five unrelated inbred *T. urticae* strains with no known history of long-term propagation on maize. In a completely randomized design that included the four maize lines, the five inbred *T. urticae* strains, and the outbred *T. urticae* W-GR strain for reference, we observed high resistance for B49, B75, and B96 to every *T. urticae* strain tested (Fig. 3). As assessed on a per mite strain basis, all analyses were significant (ANOVA, each *p* < 0.01), with B49, B75, and B96 always significantly more resistant to *T. urticae* than B73 (t-tests, *p* < 0.05, Hochberg correction for multiple testing). As assessed with each of the six mite strains, no significant differences in resistance were observed between B49, B75, and B96.

**Fig. 3.**
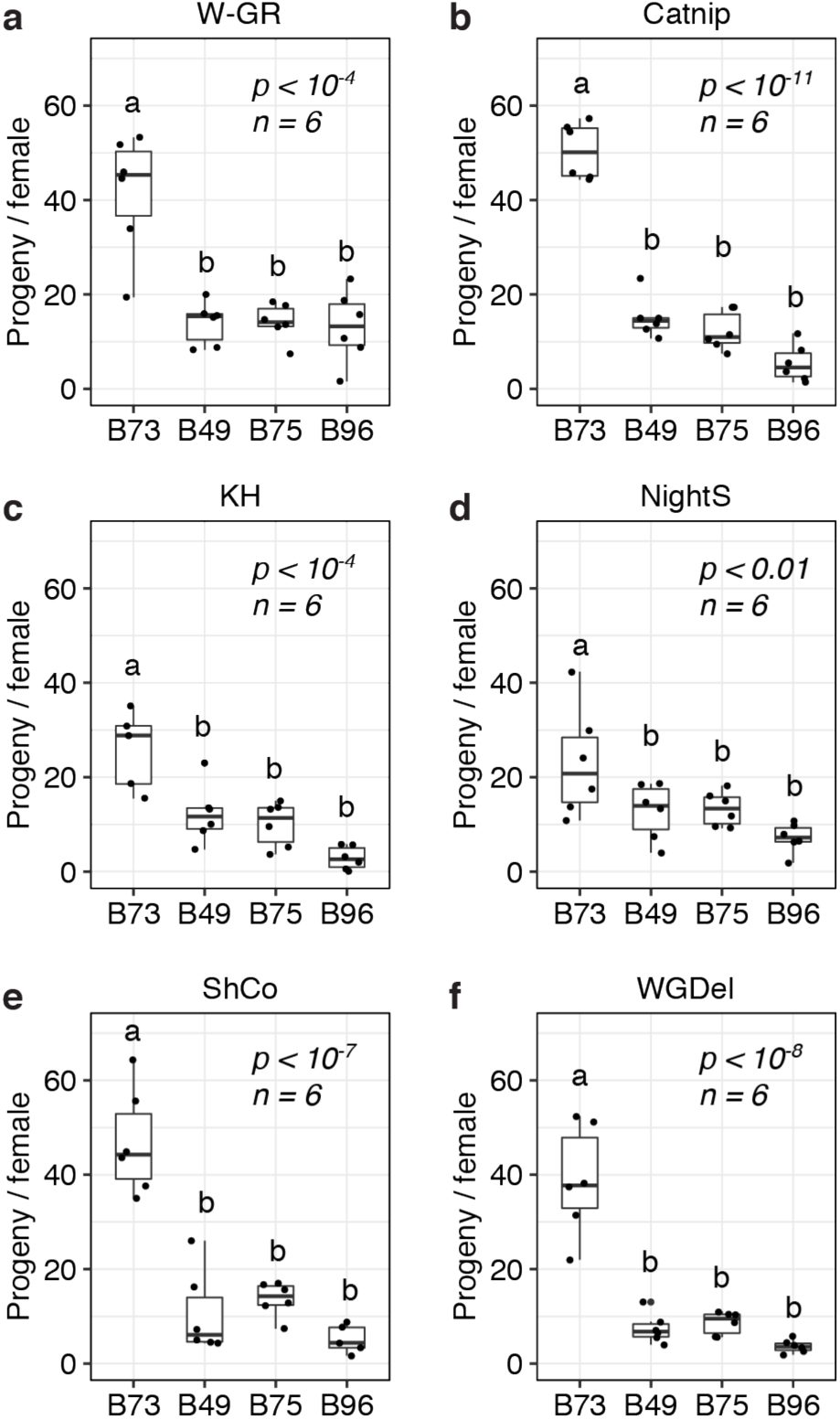
B49, B75, and B96 are resistant to multiple strains of *T. urticae*. The number of progeny per adult *T. urticae* female after six days of feeding is given for each of six unrelated *T. urticae* strains for each of four maize lines (**a**-**f**) (boxplots with overlay of data points are shown; each plot is titled by the name of the *T. urticae* strain tested). Assays were performed on the 8^th^ leaves of maize plants. Where significant variation was observed by ANOVA by mite strain (*p* < 0.05), pairwise significance was assessed with correction for multiple testing using the Hochberg method (different letters denote significant differences at *p* < 0.05). Sample size (*n*) is as indicated, except for the B73/KH (**c**) and B96/ShCo (**e**) comparisons for which one less enclosure was successfully established (*n* = 5).

### Genetic basis of *T. urticae* resistance in lines B49, B75, and B96

To identify genomic regions in maize associated with *T. urticae* resistance, we performed BSA genetic mapping with F2 populations derived from crosses of resistant B49, B75, and B96 to susceptible B73. For each F2 population, screening with *T. urticae* strain W-GR was performed with two replicates of 200 plants each, with pools of extremes of resistance (most and least resistant plants) consisting of 50 individuals each. For phenotyping, we employed a visual scale (1-7, most resistant to least resistant, respectively, as assessed with plant tissue within leaf enclosures; see Fig. S1a, and Material and Methods). Phenotyping of plant damage, as opposed to scoring mite progeny, was employed as counting mite progeny from 1200 enclosures was not practical. Nevertheless, as assessed with a subset of plants from a B49 × B73 cross for which we also counted progeny, the visual plant damage scale reflected mite productivity (*R*^*2*^=0.83, *p* < 10^−15^ Fig. S1b), the measure of resistance by antibiosis employed to identify the *T. urticae* resistance lines (Figs. 1-3).

With combined data from both replicates by cross, and using the QTLseqr package, we detected a QTL for *T. urticae* resistance on chromosome 6 centered at ∼31-86 Mb in all three crosses (for QTL detection, an FDR of 0.01 was applied to G’, a measure of genetic differentiation assessed between pools using high-throughput, short-read sequencing data, see Materials and Methods; Fig. 4). For the B49 × B73 and B75 × B73 crosses, this QTL was the only major QTL detected. In contrast, for B96, a second QTL with similar G’ peak values was also apparent for a broader peak region extending from ∼29-167 Mb on chromosome 1 (Fig. 4c). In all of these cases, alleles contributed by the *T. urticae* resistant maize lines were enriched in the resistant BSA pools in the QTL intervals. When we performed the same analyses using each replicate for each cross individually, we obtained the same or similar results (i.e., for each independent replicate in each cross, the chromosome 6 region was significant as a QTL, and in the B96 × B73 cross, the region on proximal chromosome 1 was also identified as a QTL in each independent replicate; Fig. S2-4). In several cases, in either the analyses with combined replicates, or in the analyses with individual replicates, G’ values reached or nearly reached the QTL detection threshold elsewhere in the genome. For example, on chromosome 5 for both the B75 × B73 and B96 × B73 crosses as assessed with both replicates (Fig. 4b,c), or for one replicate of the B96 × B73 cross but not the other (Fig. S4). Regardless, as assessed with both replicates (Fig. 4), the magnitude of all G’ peaks, except those on chromosomes 1 and 6, was minor (see also Discussion).

**Fig. 4.**
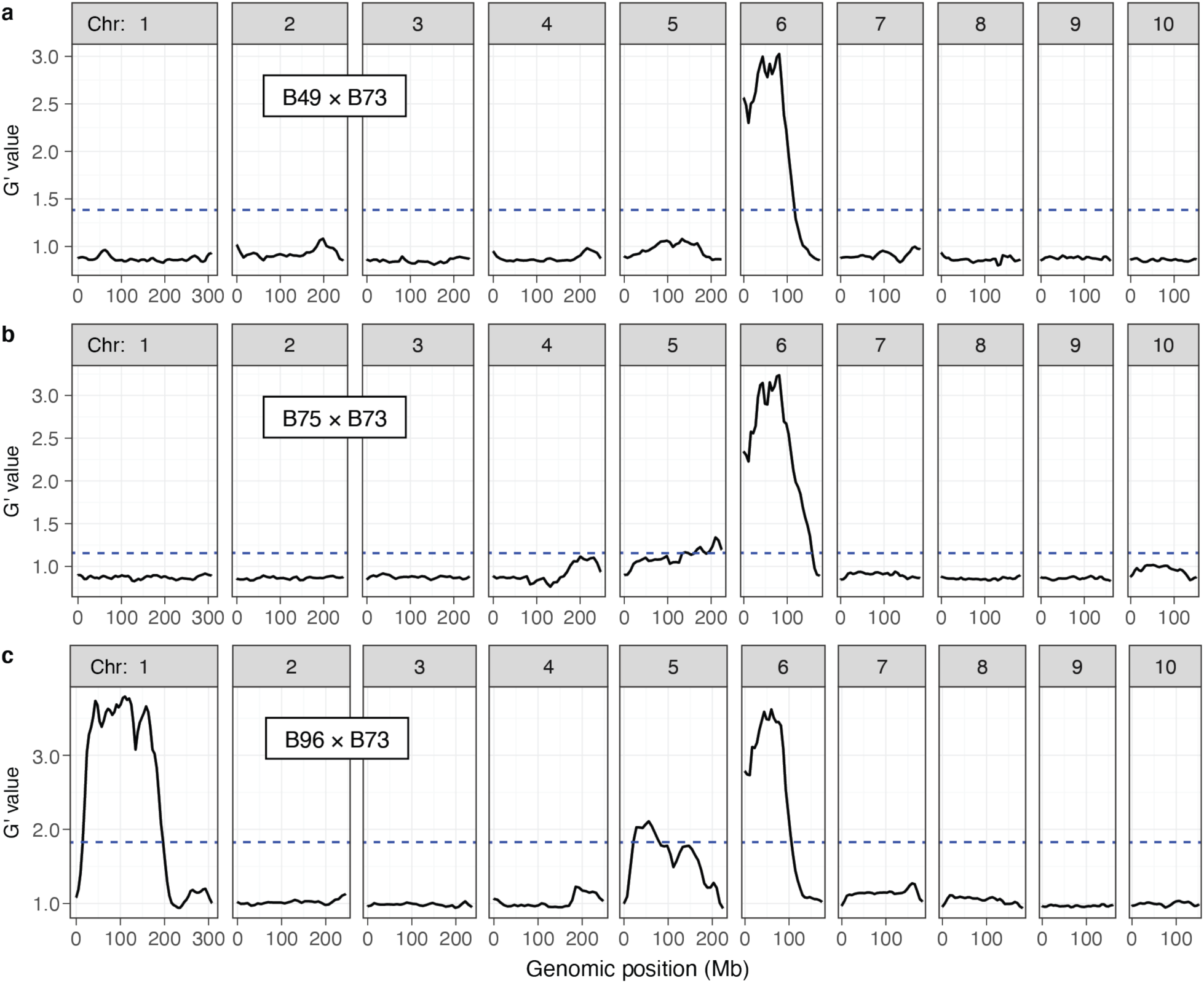
Resistance QTL to *T. urticae* as identified by BSA genetic mapping in F2 maize populations. Shown are plots of G’ obtained from contrasting pools of *T. urticae* strain W-GR resistant and sensitive plants from crosses of resistant maize lines B49, B75 and B96 to susceptible line B73 (**a, b**, and **c**, respectively; chromosomes 1-10 are as indicated, left to right). Deflections in G’ values exceeding a genome-wide false discovery rate of 0.01 (dashed lines) identify QTL. Extent of resistance was based on leaf damage scores on the 4^th^ leaves, and for each cross the panels represent the combined analysis of two experimental replicates (combined, 100 plants in each of the resistant and susceptible pools of phenotypic extremes; see Materials and Methods). Respective individual analyses for each of the two replicates by cross, B49 × B73, B75 × B73, and B96 × B73, are given in Fig. S2, Fig. S3, and Fig. S4, respectively. The plots shown were modified from the output of QTLseqr (see Materials and methods).

As additional confirmation of QTL detection by the BSA method, for the B49 × B73 and B75 × B73 crosses we performed single locus genotyping with a PCR marker centered on the chromosome 6 QTL interval. As assessed by genotyping all F2 individuals in a replicate for cross B49 x B73 and a replicate for cross B75 x B73, resistance scores were significantly different among the genotypic classes (ANOVA, each *p* < 10^−11^; Fig. 5). Further, in each case resistance appeared to be recessive (or nearly so) as plants homozygous for the resistant parent at the chromosome 6 marker were significantly different from plants either homozygous or heterozygous for the B73 allele (t-tests, *p* < 0.05, Hochberg correction for multiple testing), with the latter two genotypic classes not significantly different from each other. As assessed with the homozygous classes, 70.6% and 61.2% of the variances in resistance were explained by the marker genotype in the chromosome 6 QTL region in the B49 × B73 and B75 × B73 crosses, respectively (*R*^*2*^, *p* < 10^−15^ in each case). In particular, the distributions of damage scores for the homozygous classes for the B49 × B73 cross were almost non-overlapping, excepting for a prominent outlier that plausibly reflects recombination between the marker and the causal resistance locus (Fig. 5a, homozygous B49 bin, upper-most data point). Excluding this outlier, 80.1% of the variance in resistance was explained by the chromosome 6 marker genotype in the B49 × B73 cross.

**Fig. 5.**
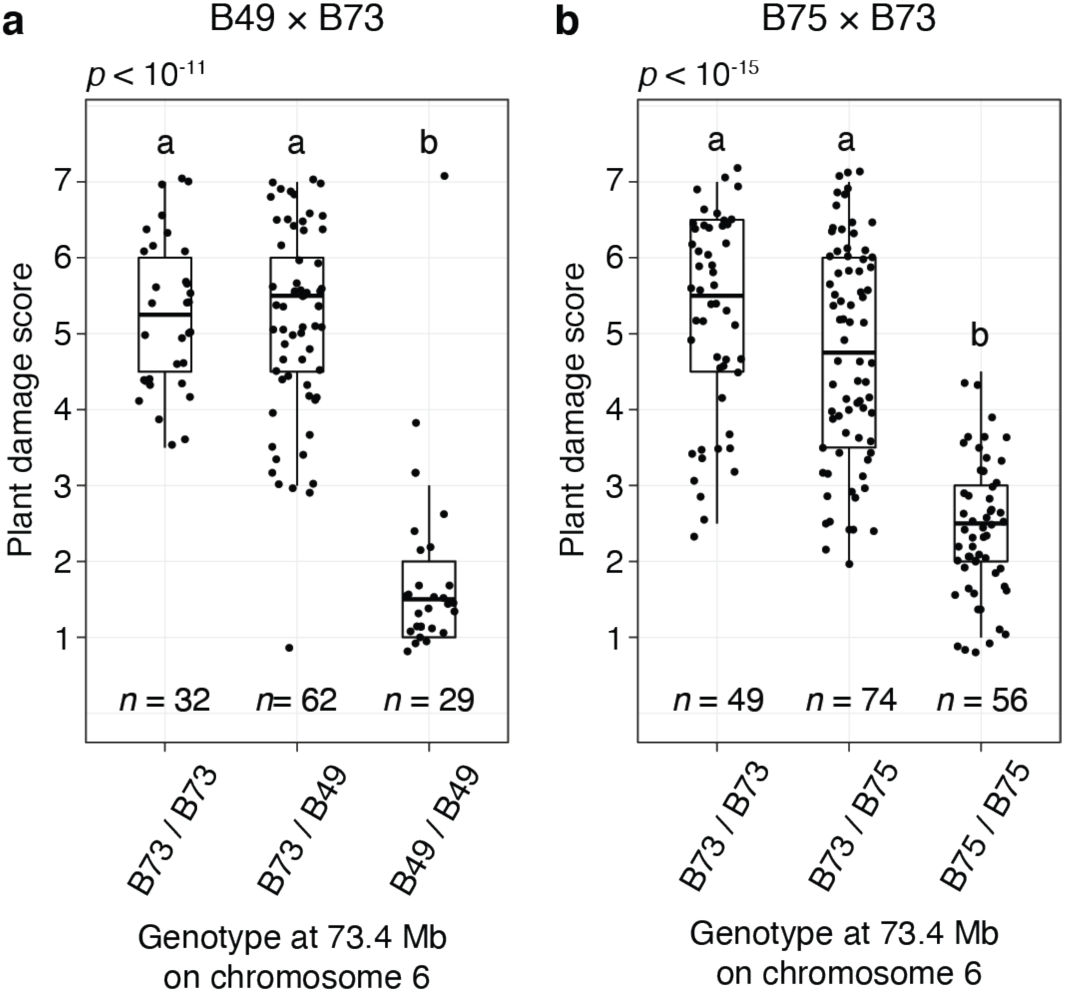
The chromosome 6 *T. urticae* resistance QTL has a recessive mode of action in B49 and B75 as assessed in crosses to susceptible B73. Shown are boxplots with an overlay of data points for resistance to *T. urticae* strain W-GR as assessed with a plant damage score and classified by genotype at a locus at 73.4 Mb on chromosome 6 that is within the peak chromosome 6 QTL interval (Fig. 4). The data shown are from BSA replicate two of cross B49 × B73 (**a**) and BSA replicate one of cross B75 × B73 (**b**). Data for all plants for which genotyping was successful in each replicate population are shown (i.e., not just the plants comprising the pools of phenotypic extremes used in BSA scans, Fig. 4, Fig. S2b, and Fig. S3a). Where significant variation was observed by ANOVA by cross (*p* < 0.05), pairwise significance was assessed with correction for multiple testing using the Hochberg method (different letters denote significant differences at *p* < 0.05).

### A shared basis for resistance among B49, B75 and B96

B49 was derived from a cross with B96 as one parent, while B75 was derived in a breeding program involving 16 parents, including B49 (Portwood et al. 2019). The shared ancestry of the three *T. urticae* resistant lines, the coincident location of a large-effect resistance QTL on chromosome 6, as well as its shared mode of action in crosses of B73 to both B49 and B75 (recessive resistance), raised the possibility of a common genetic basis for *T. urticae* resistance among the three lines. To assess this possibility, we sequenced the genomes of B49, B75, and B96, and used resulting high-quality SNP predictions to identify genomic intervals of identity (recent ancestry) in pairwise comparisons as well as across all three lines (Fig. 6a and 6b, respectively). As revealed from sliding window analyses, and as expected from the B49 pedigree, many large regions of the B49 genome were identical to B96 (Fig. 6a). In contrast, only several extended chromosome regions from B75, which is more distantly related to B96, showed a similar pattern. However, much of chromosome 6 was identical between B49, B75, and B96, including for the region harboring the chromosome 6 resistance QTL (compare Fig. 6 to Fig. 4). In contrast, no extended regions of identity were observed between B96 and either B49 or B75 on chromosome 1 in the interval for the resistance QTL that is unique to B96.

**Fig. 6.**
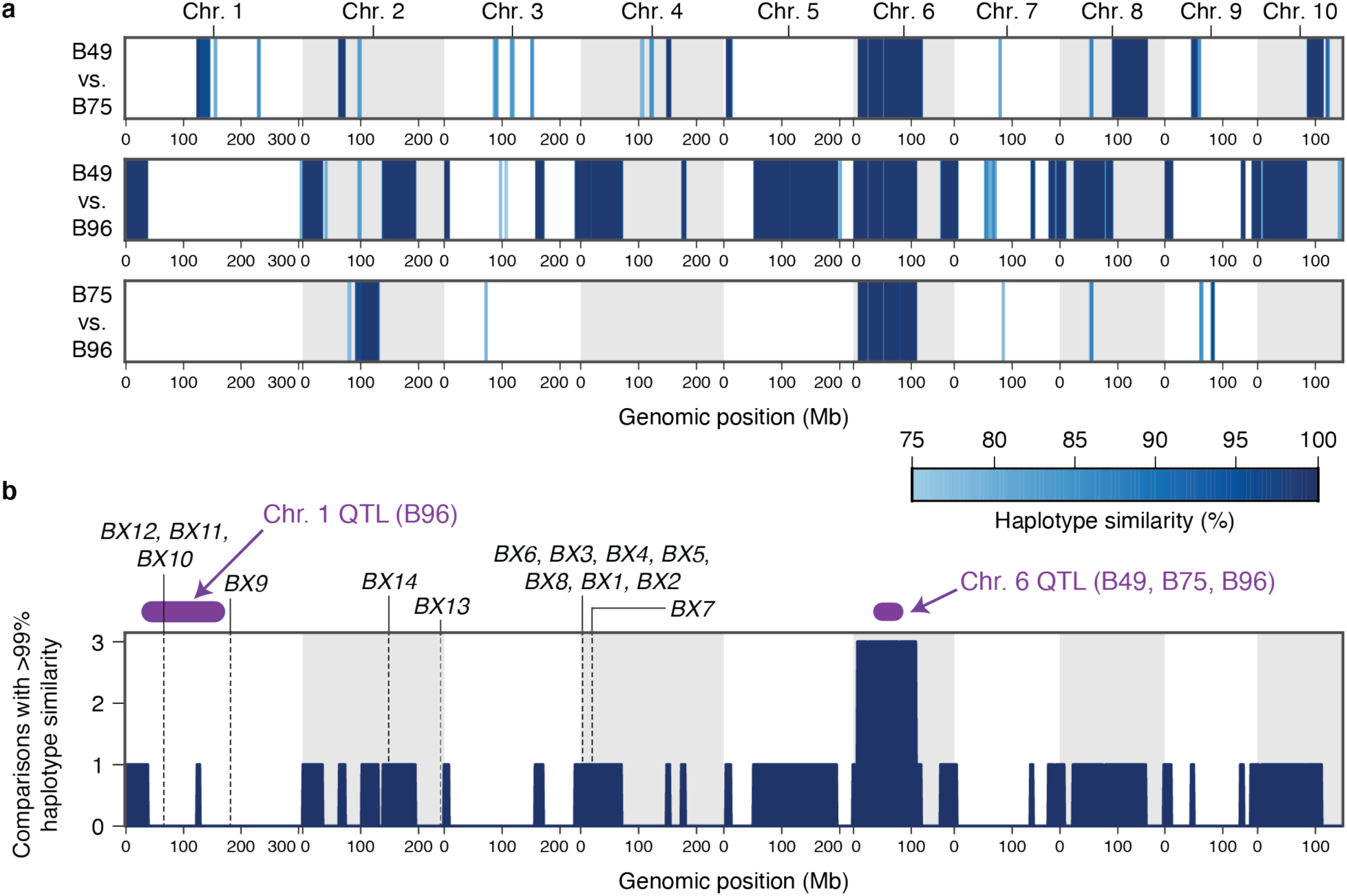
A single haplotype on chromosome 6 associates with high-level resistance to *T. urticae* in maize lines B49, B75 and B96. (**a**) Haplotype similarities as assessed with SNP data in sliding window analyses are shown for all pairwise comparisons between B49, B75 and B96. The depth of blue shading indicates the % haplotype similarity (see scale, lower right). (**b**) Genomic regions for which the identical (or nearly identical) haplotypes are observed in multiple comparisons among the three resistance maize lines (comparisons with >99% similarity, dark blue). The location of peak genomic intervals for major QTL identified on chromosome 1 (B96) and chromosome 6 (B49, B75, and B96) (Fig. 4 and Results) are in purple as indicated, and the location of known benzoxazinoid biosynthetic genes (or gene clusters) are indicated by vertical dashed lines. In both panels, chromosomes are demarcated by alternating white and gray backgrounds.

### Benzoxazinoid biosynthesis genes and QTL candidate intervals

As expected for genetic mapping with F2 populations, the approximate QTL peak regions for *T. urticae* resistance on chromosomes 1 and 6 include large genomic intervals. As assessed from peaks of G’ in the BSA analyses, the chromosome 1 and chromosome 6 peak QTL regions that extend from ∼29-167 Mb (138 Mb) and ∼31-86 Mb (55 Mb) (Fig. 4 and Fig. 6b) include 2,645 and 567 annotated protein coding genes, respectively. To date, the only endogenous maize plant defenses implicated in antibiosis to *T. urticae* are benzoxazinoids (Bui et al. 2018). While no genes encoding benzoxazinoid biosynthetic enzymes are located on chromosome 6, the *BX10, BX11* and *BX12* biosynthetic gene cluster is located at ∼66.3 Mb on chromosome 1, and is therefore within the peak interval for the B96 chromosome 1 QTL (Fig. 6b). Another benzoxazinoid biosynthetic gene, *BX9*, which is located at 182.3 Mb on chromosome 1, is nearby but distal to the peak chromosome 1 QTL region.

## Discussion

Previous work has suggested that while most maize lines are comparatively resistant to *T. urticae* at the seedling stage, they subsequently become susceptible (Mansour et al. 1993; Tadmor et al. 1999). This inversely correlates with constitutive benzoxazinoid levels, which impact *T. urticae* (Bui et al. 2018), are high in seedlings of many lines, and begin to decrease rapidly by the ∼2-3 leaf stage (Bing et al. 1990b), reflecting changes in defensive programs as maize plants mature. Nevertheless, moderate-to high-level resistance in some maize lines to both *T. urticae* and *O. pratensis* herbivory has been reported in studies that included plants at older stages (Kamali et al. 1989; Mansour et al. 1993; Tadmor et al. 1999; Bynum et al. 2004b), an important consideration as spider mites invade fields throughout the growing season to cause damage of economic importance on more mature plants (Peairs 2010). In our study, we assessed reproduction of both *T. urticae* and *O. pratensis* on 38 maize lines at an agriculturally relevant post-seedling stage (the 8^th^ leaf). Our findings were broadly consistent with some, but not all, earlier studies.

As reported in previous screens assessing antibiosis, we found that most maize lines were comparatively susceptible to *T. urticae*, but identified a small number of lines, including B49, B75, and B96, as highly resistant to the W-GR *T. urticae* strain. Our results confirm that B96 exhibits high antibiosis to *T. urticae*, as observed previously in both laboratory and field settings at post-seedling stages, and with unrelated mite strains (Kamali et al. 1989; Tadmor et al. 1999). Further, B49 was reported previously to be resistant to *T. urticae* at the seedling stage (Tadmor et al. 1999); to our knowledge, B75 has not been screened previously for spider mite resistance. While Oh43, a NAM line founder, was reported to be highly resistant to *T. urticae* as assessed on detached leaf segments at the 4^th^-leaf stage (Mansour, F. and Bar-Zur, A. 1992), it was as susceptible as the majority of the other 37 lines screened in the current study. Our observation that Oh43 was not highly resistant may be explained by our use of older plants for screening, consistent with the finding of Kamali et al. (1989) who observed only modest resistance for Oh43 in a field trial using older plants. Alternatively, it might reflect the use of different *T. urticae* strains, as intra-specific variation in fitness on different plant hosts has been reported for field-derived or laboratory-evolved *T. urticae* strains (Wybouw et al. 2019; Sousa et al. 2019). The three lines we characterized as highly resistant to the *T. urticae* W-GR strain were, however, also resistant to five unrelated *T. urticae* strains with no known, recent history of propagation on maize. Therefore, resistance loci and alleles from B49, B75 and B96 are likely to be generally applicable to efforts to develop *T. urticae* resistant maize varieties.

Despite their resistance to *T. urticae*, neither B49, B75, B96, nor any of the other 35 maize lines we studied, exhibited marked antibiosis to the *O. pratensis* strain used in our study. This contrasts with the finding for *T. urticae*, but is consistent with Bui et al.’s (2018) work suggesting that the generalist *T. urticae* and the grass specialist *O. pratensis* respond differently to maize defenses, as well as with the plausible expectation that the grass-specialist *O. pratensis* may have evolved more robust mechanisms to overcome the defenses of its host than the generalist herbivore. Additionally, it should be noted that although resistance by antibiosis to *O. pratensis* has been reported for some maize lines, in a number of cases resistance arising from tolerance was demonstrated or is plausible (Owens et al. 1976; Bynum et al. 2004b). We included the founders of the maize NAM population in our study as this genetic resource has facilitated the identification of QTL for resistance to insect herbivores (Meihls et al. 2013; Tzin et al. 2015). However, as our most *T. urticae* resistant maize lines were not NAM founders, we instead exploited BSA genetic mapping to identify resistance QTL in B49, B75, and B96. It should be noted that given the scope of the BSA screens, of necessity we used younger plants, and we used a plant damage metric to assess resistance (however, as assessed with one cross, plant damage was highly correlated with mite reproduction, suggesting that our BSA scans assessed resistance arising from antibiosis; Fig S1b). Each of the three lines harbored a large-effect QTL on chromosome 6, which as assessed by single-locus genotyping in B49 and B75 crosses conferred recessive resistance, and a second resistance QTL on chromosome 1 was unique to B96. In B75 and B96, a possible minor-effect locus or loci, or ones that are environmentally sensitive, may be located on chromosome 5 (i.e., chromosome 5 was identified as a QTL in only one of the two BSA replicates for the B96 × B73 F2 mapping population, which were performed at different times albeit in the same greenhouse bays, Fig. S4, and Materials and Methods). Nonetheless, a possible role for chromosome 5 in resistance requires further study. As revealed from haplotype analyses using genomic sequence data, the shared chromosome 6 QTL was underpinned by an identical haplotype putatively derived from B96, a parent of both B49 and B75 (Portwood et al. 2019). Thus, a single haplotype putatively originating from B96 underlies high-level resistance to *T. urticae* in our sizeable study population of highly diverse maize lines. The identification of a second chromosome 1 resistance QTL in B96 would seem to be at odds with the observation that we did not observe statistically significant increases in resistance for B96 as compared to either B49 or B75. Nevertheless, for nearly all cases where we compared resistance between the three lines, B96 had the lowest median number of *T. urticae* progeny (Fig. 1a, Fig. 2a, and Fig. 3), consistent with the detection of an additional resistance locus.

B96 has been reported to be “nearly immune” to first-generation feeding by the European corn borer, *O. nubilalis* (Kamali et al. 1989), and was also observed to be highly resistant to a complex of three thrips species (Bing et al. 1990a). While it has poor agronomic properties, B96 was used to produce *O. nubilalis* resistant B49, B64, and B68, which were used in additional breeding programs, including for the generation of B75 that is also resistant to *O. nubilalis* (Portwood et al. 2019). B96 has also been reported to have high levels of DIMBOA at late developmental stages (Bing et al. 1990b), as has B75 (Barry et al. 1994). The observation that *T. urticae* reproduction is inhibited by benzoxazinoids (Bui et al. 2018), and that the *T. urticae* resistant lines in the current study are resistant to *O. nubilalis*, for which benzoxazinoids are also detrimental (Wouters et al. 2016), raises the possibility of a shared molecular-genetic basis of resistance. However, such conjecture must be interpreted with caution. For example, B64 and B68 were included in our study and were comparatively susceptible to *T. urticae*, as was line CI31A, which is both resistant to *O. nubilalis* and is a high DIMBOA content line (Barry et al. 1994; Portwood et al. 2019). One interpretation of these findings is that molecular-genetic resistance mechanism(s), if shared between *T. urticae* and insects, may only partially overlap. An additional confounding factor is that while DIMBOA content has been reported for some lines in our study, the levels of other benzoxazinoids have not. For instance, the production of the DIMBOA derivative 2-hydroxy-4,7-dimethoxy-1,4-benzoxazin-3-one (HDMBOA) is thought to be especially important for defense against lepidopteran pests (Glauser et al. 2011), and absolute and relative levels of different benzoxazinoid metabolites can vary substantially among maize lines (Meihls et al. 2013). Supporting a possible role for benzoxazinoids in explaining intraspecific maize variation for *T. urticae* resistance comes from the observation that the chromosome 1 QTL region in B96 harbors benzoxazinoid biosynthetic genes in the *BX10-12* gene cluster involved in the synthesis of HDMBOA (Meihls et al. 2013). Moreover, although no known benzoxazinoid biosynthetic genes are located on chromosome 6, the peak region for the chromosome 6 QTL common to B49, B75, and B96 (∼31-86 Mb) harbors the putative transcriptional regulator *NACTF21* (at 69.0 Mb on chromosome 6). Recently, *NACTF21* was implicated in a large-scale transcriptome network analysis as a potential regulator of benzoxazinoid biosynthetic genes, including *BX1* and *BX2* (Zhou et al. 2020), whose products perform the first two enzymatic steps required for the synthesis of all benzoxazinoids (Wouters et al. 2016). However, while benzoxazinoid biosynthetic genes and putative *trans* regulators should be investigated as potentially causal for the chromosome 1 and 6 QTLs, the peak regions identified in the BSA scans contain many genes. This is especially true for the broad QTL region identified on chromosome 1 in B96, and the involvement of multiple, linked resistance loci cannot be ruled out.

In conclusion, we show that the genetic basis of variation in maize resistance to the generalist *T. urticae* and the specialist *O. pratensis* differ, and we identified two QTL underlying high-level resistance to *T. urticae*, the most geographically widespread spider mite pest of maize (Migeon et al. 2010). The loci we identified should be of use in marker-assisted breeding programs to develop maize lines resistant to this generalist mite. Moreover, they serve as entry points for future studies to identify the specific molecular-genetic underpinnings of maize resistance to *T. urticae*, as well as to investigate if intra-specific variation in maize resistance pathways to spider mites also impacts that of major insect pests.

## Acknowledgements

This work was supported by USA National Science Foundation Plant Genome Research Program award 1444449 to R.M.C and R.A.R. Additionally, the generation of the inbred mite lines was supported by USA National Science Foundation award 1457346 to R.M.C.

## Statement of Data Availability

Illumina read data for maize inbred lines B49, B75, and B96, and for BSA pools derived from crosses of these lines to B73, has been deposited to the USA National Center for Biotechnology Information (NCBI) Sequence Read Archive (SRA) under PRJNA481365 and PRJNA556665. Phenotypic data, and genotypic data as VCF files, for lines B49, B75, and B96, and for BSA pools derived from crosses of these lines to B73, are publicly available as datasets on the figshare repository (https://doi.org/10.6084/m9.figshare.13708375.v1).

## Author contributions statement

HB, RMC, GSG, and RAR originated the study and experimental design. HB, RMC, GSG, CR, and SL performed plant-mite interaction experiments. AK, HB, and RMC created mite inbred lines. HB, RG, AHK, and MJ performed bioinformatic and statistical analyses. HB and RMC assumed the primary role in writing the manuscript, which was reviewed and approved by all authors.

## Supplementary Figure Legends

**Fig. S1** Relationship between plant damage and mite reproduction in a cross between the *T. urticae* resistant maize line B49 and susceptible B73. (**a**) An image for a representative highly resistant sample (score 1) and a representative highly susceptible sample (score 7) following six days of *T. urticae* feeding (4^th^ leaves are displayed). Damage to the lower (abaxial) and upper (adaxial) leaf blades are shown; the leaf segments were excised from within Tanglefoot enclosures for image capture. For the resistant sample (right), a region of reduced feeding (outlined by dashed white lines) corresponds to where tape was affixed to apply the pipet tip of mites for release into the leaf blade enclosure (see Materials and Methods). Scale bars: 1 cm. (**b**) Boxplots (plant damage scores, plotted in units of 0.5, with jitter of data points by bin) showing the relationship between the visual plant damage score and mite productivity – as assessed by the number of progeny counted following six days of addition of adult *T. urticae* females to enclosures – for a subset of plants used for BSA genetic mapping (data shown are for replicate two of the B49 cross, Fig. S2b). Blue circles represent individual data points.

**Fig. S2** BSA genetic mapping scans for QTL for resistance to *T. urticae* strain W-GR in two F2 population replicates of the B49 × B73 cross. The plots shown correspond to that of Fig. 4a, except that the BSA analysis was done on a per replicate basis as indicated, top and bottom (see the legend for Fig. 4 for additional details). The solid red line denotes a genome-wide FDR of 0.01 for QTL detection.

**Fig. S3** BSA genetic mapping scans for QTL for resistance to *T. urticae* strain W-GR in two F2 population replicates of the B75 × B73 cross. The plots shown correspond to that of Fig. 4b, except that the BSA analysis was done on a per replicate basis as indicated, top and bottom (see the legend for Fig. 4 for additional details). The solid red line denotes a genome-wide FDR of 0.01 for QTL detection.

**Fig. S4** BSA genetic mapping scans for QTL for resistance to *T. urticae* strain W-GR in two F2 population replicates of the B96 × B73 cross. The plots shown correspond to that of Fig. 4c, except that the BSA analysis was done on a per replicate basis as indicated, top and bottom (see the legend for Fig. 4 for additional details). The solid red line denotes a genome-wide FDR of 0.01 for QTL detection.

