## Supplementary figures and images for "Maize inbred line B96 is the source of large-effect loci for resistance to generalist but not specialist spider mites"

### Supplemental Figure 1

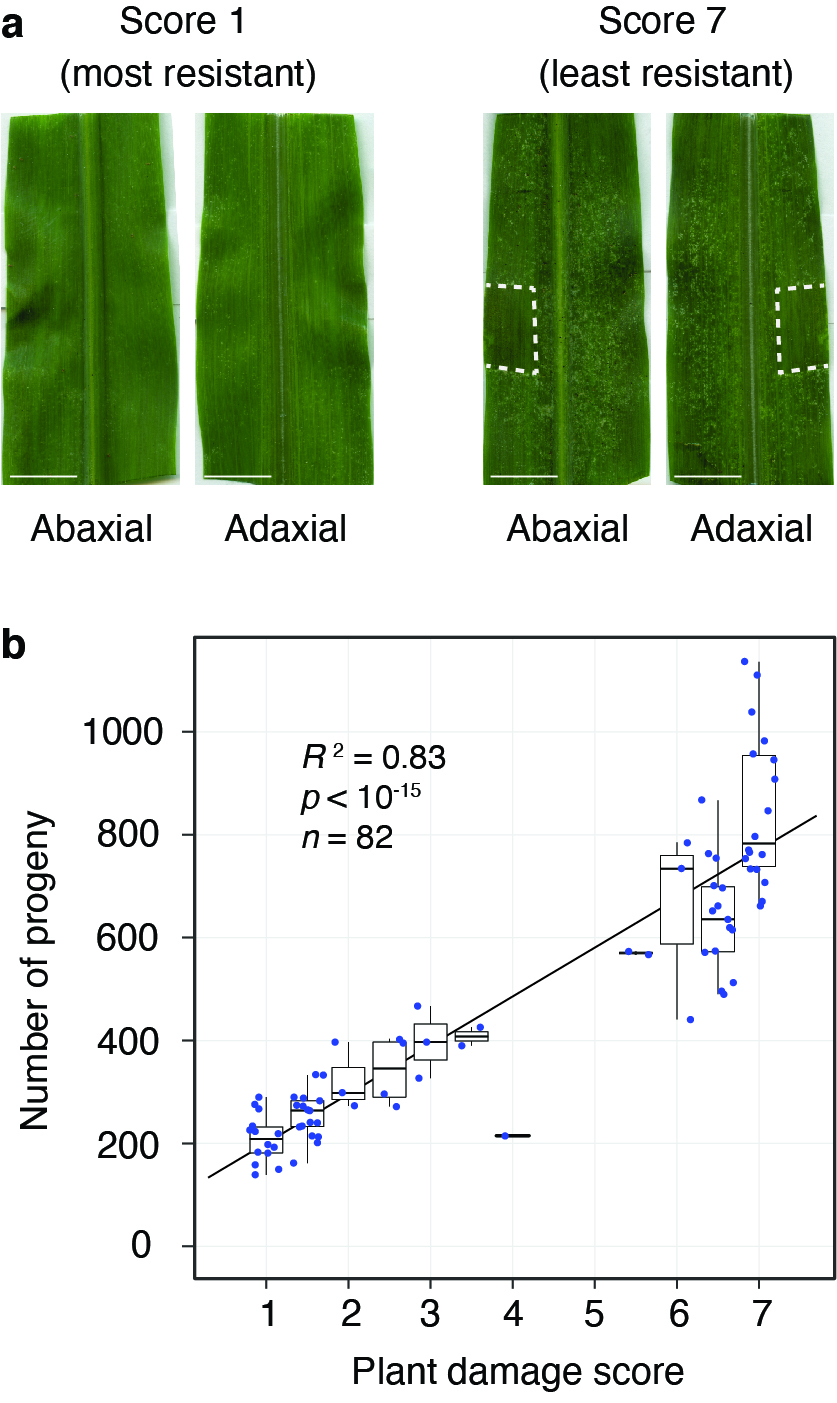

### Supplemental Figure 2

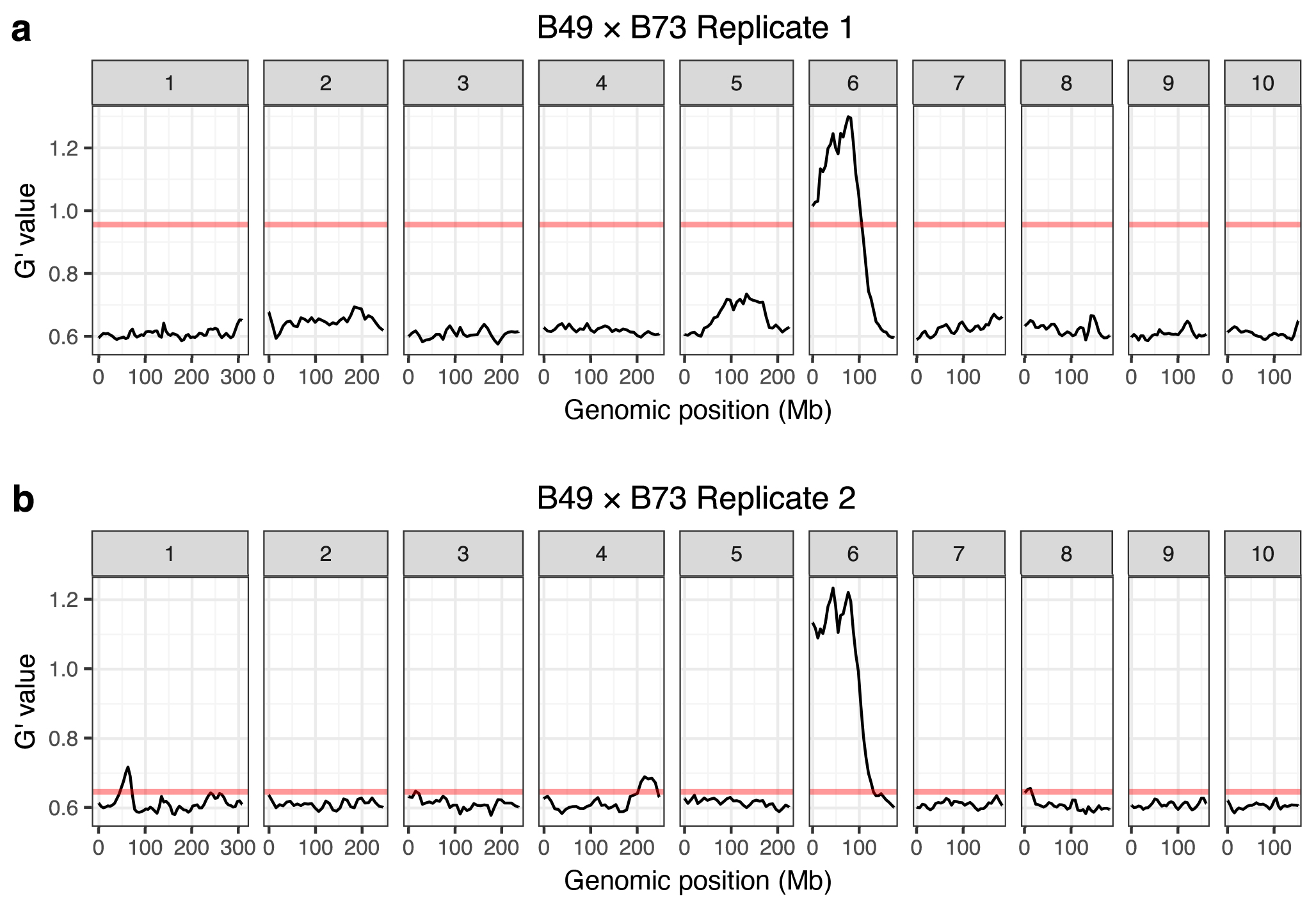

### Supplemental Figure 3

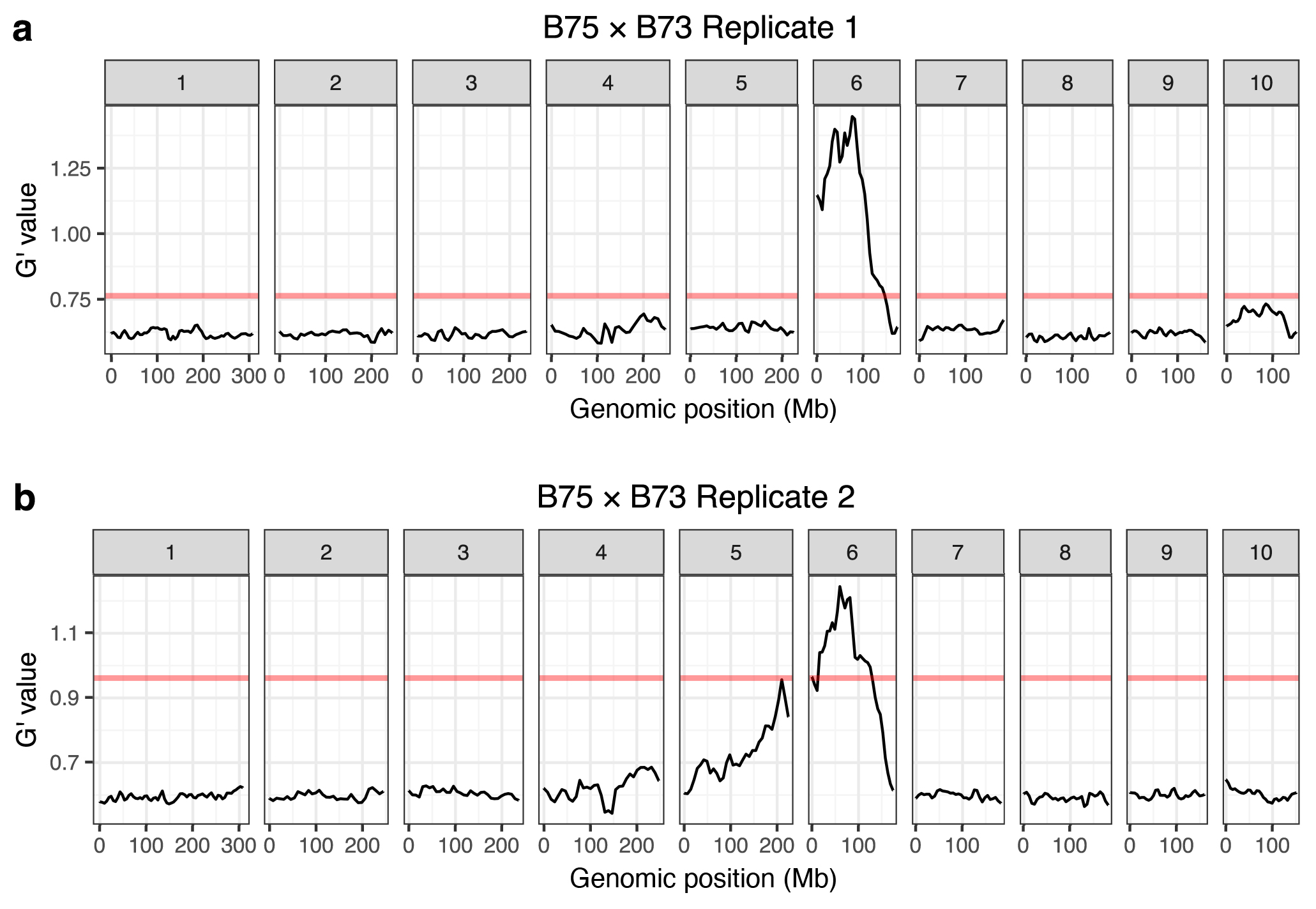

### Supplemental Figure 4

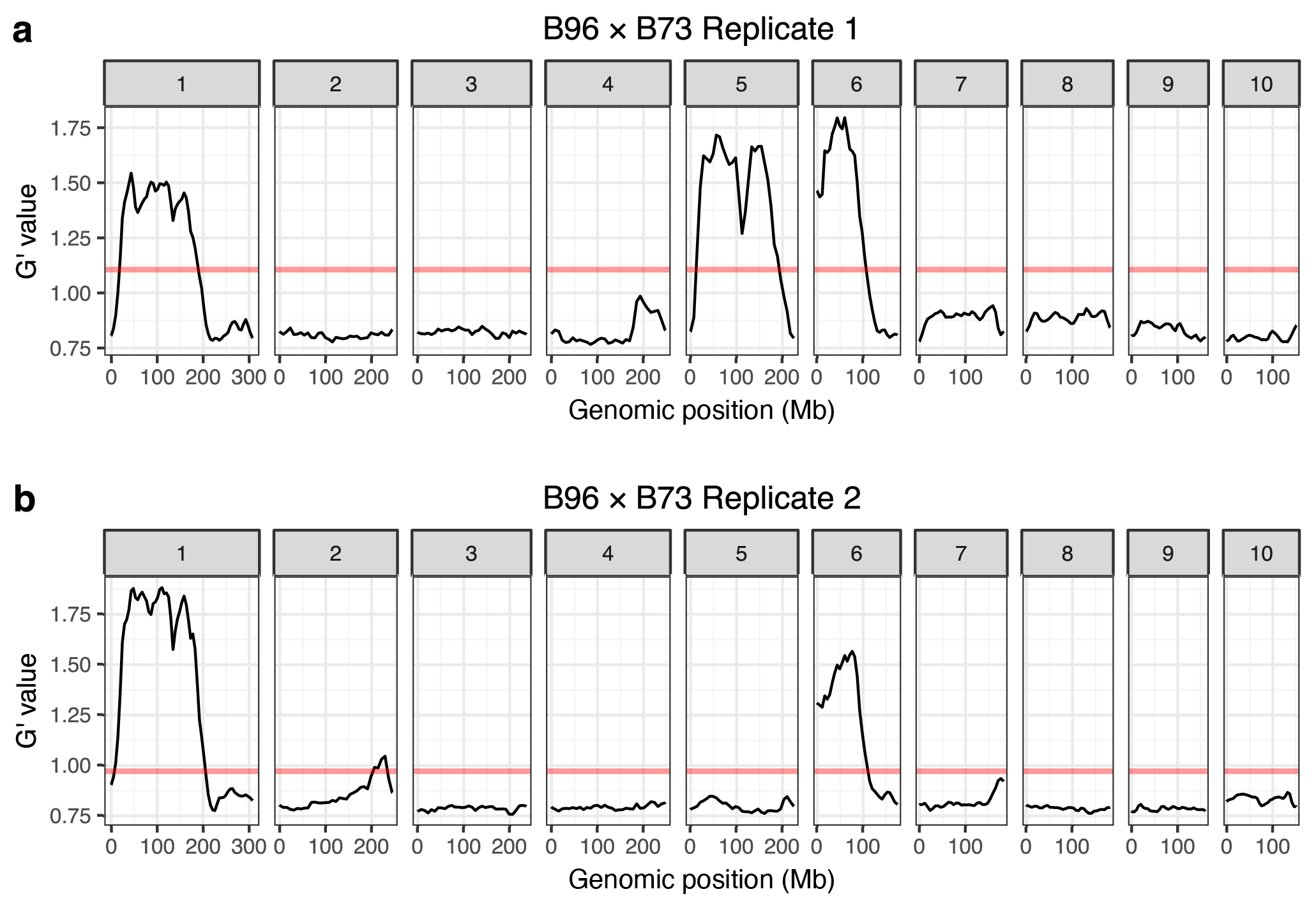
